# Dietary cysteine enhances intestinal stemness via CD8^+^ T cell-derived IL-22

**DOI:** 10.1101/2025.02.15.638423

**Authors:** Fangtao Chi, Qiming Zhang, Jessica E.S. Shay, Johanna Ten Hoeve, Yin Yuan, Zhenning Yang, Heaji Shin, Sumeet Solanki, Yatrik M. Shah, Judith Agudo, Ömer H. Yilmaz

## Abstract

A critical question in physiology is understanding how tissues adapt and alter their cellular composition in response to dietary cues. The mammalian small intestine, a vital digestive organ that absorbs nutrients, is maintained by rapidly renewing Lgr5^+^ intestinal stem cells (ISCs) at the intestinal crypt base. While Lgr5^+^ ISCs drive intestinal adaptation by altering self-renewal and differentiation divisions in response to diverse diets such as high-fat diets and fasting regimens, little is known about how micronutrients, particularly amino acids, instruct Lgr5^+^ ISC fate decisions to control intestinal homeostasis and repair after injury. Here, we demonstrate that cysteine, an essential amino acid, enhances the ability of Lgr5^+^ ISCs to repair intestinal injury. Mechanistically, the effects of cysteine on ISC-driven repair are mediated by elevated IL-22 from intraepithelial CD8αβ^+^ T cells. These findings highlight how coupled cysteine metabolism between ISCs and CD8^+^ T cells augments intestinal stemness, providing a dietary approach that exploits ISC and immune cell crosstalk for ameliorating intestinal damage.

## Introduction

The mammalian intestine plays a central role in the digestion and absorption of nutrients, including carbohydrates, amino acids, and lipids. Rapidly renewing Lgr5^+^ intestinal stem cells (ISCs) domiciled at the intestinal crypt base actuate intestinal epithelial adaptation to diverse diets by regulating the balance between self-renewal and differentiation divisions. For example, in response to long-term calorie restriction, dampened mTOR complex 1 (mTORC1) signaling in the Paneth cell niche boosts ISC numbers and their ability to repair intestinal injury^1^. Ad libitum refeeding after a 24-hour fast cell-autonomously augments ISC-driven regeneration by activating an mTOR-polyamine metabolite-protein translation axis^2^. Finally, high-fat diets or dietary lipids such as fatty acids and cholesterol species augment ISC activity by activating peroxisome proliferator-activated receptor (PPAR) programs or phospholipid remodeling^3,4,5,6,7,8^. Although macronutritional changes such as fasting/refeeding or high-fat diets alter ISC fate decisions and activity, little is known about how specific amino acids impact ISC fate decisions or regenerative capacity.

The intestinal stem cell (ISC) niche comprises a complex interplay of epithelial Paneth cells, specialized stromal fibroblasts, neurons, and immune cells. This microenvironment supports Lgr5^+^ ISCs by producing crucial factors, including WNTs, RSPOs, BMP inhibitors, and cytokines^9,10,11,12,13^. Mucosal immune cells, such as T cells, innate lymphoid cells (ILCs), dendritic cells, and macrophages, mediate host defenses and inflammatory responses and directly modulate ISC function^14^. In particular, CD8^+^ T cells, comprised of regulatory αα^+^ and αβ^+^ T subsets, play a dynamic role in regulating intestinal epithelial adaption to diet^15,16,17^. CD8αα^+^ γδ T cells, for example, facilitate the induction of transporters and enzymes that permit epithelial adaptation to carbohydrate-rich diets^18^; yet, whether and how specific nutrients control ISC activity via changes in intestinal CD8αβ^+^ T cells is unknown.

IL-22 is a critical immune cell-derived cytokine that enhances ISC-mediated regeneration following injury^10,19,20,21^. Mice with IL-22 pathway deficiencies exhibit reduced Lgr5^+^ ISC frequency and impaired epithelial regeneration in response to injury and inflammation^21,22,23,24^. However, precise mechanisms by which dietary-induced metabolic changes regulate IL-22 production in specific CD8^+^ T cell populations, like CD8αβ^+^ T cells, and how this impacts ISC function remains under-explored. Our research reveals that the essential amino acid cysteine plays a previously unappreciated role in enhancing ISC function and tissue regeneration via IL-22 production in intestinal CD8αβ^+^ T cells, highlighting how coupled cysteine metabolism between ISCs and CD8αβ^+^ T cells augments intestinal stemness.

### Amino acids show various impacts on ISC function

We and others identified that HMGCS2, the rate-limiting enzyme for ketone metabolite production, is a key metabolic enzyme that defines the small intestinal Lgr5^+^ ISC gene expression signature^7,25^. Unlike other ISC markers such as LGR5 and OLFM4, HMGCS2 expression dynamically responds to dietary interventions where levels of HMGCS2 and its ketone metabolites directly correlate with intestinal stemness and regenerative capacity^3,4,5,6,7^(**Fig. 1a**). To gain insights into how amino acids influence intestinal HMGCS2 expression, we investigated how each amino acid controlled HMGCS2 expression within small intestinal crypts. Each of the 20 proteinogenic amino acids was administered orally to mice in two doses, 24 hours apart, followed by intestinal tissue collection 24 hours after the final dose for immunohistochemical (IHC) analysis. Notably, cysteine treatment significantly induced HMGCS2 expression in crypts, which is typically expressed in crypt base Lgr5+ ISCs of the small intestine (**Fig. 1b**). Phenylalanine, arginine^26^, and leucine also modestly increased HMGCS2 expression (**Fig. 1b**). However, methionine, another sulfur-containing amino acid, slightly suppressed HMGCS2 expression (**Fig. 1b**). Next, we assessed cysteine related metabolites, including glutathione (GSH), taurine, and hypotaurine, but found they do not control intestinal HMGCS2 expression (**Extended Data Fig. 1a-b**). These observations indicate that cysteine mediates its effects on intestinal HMGCS2 expression through mechanisms independent of its antioxidant properties (**Extended Data Fig. 1b)**.

**Fig. 1.**
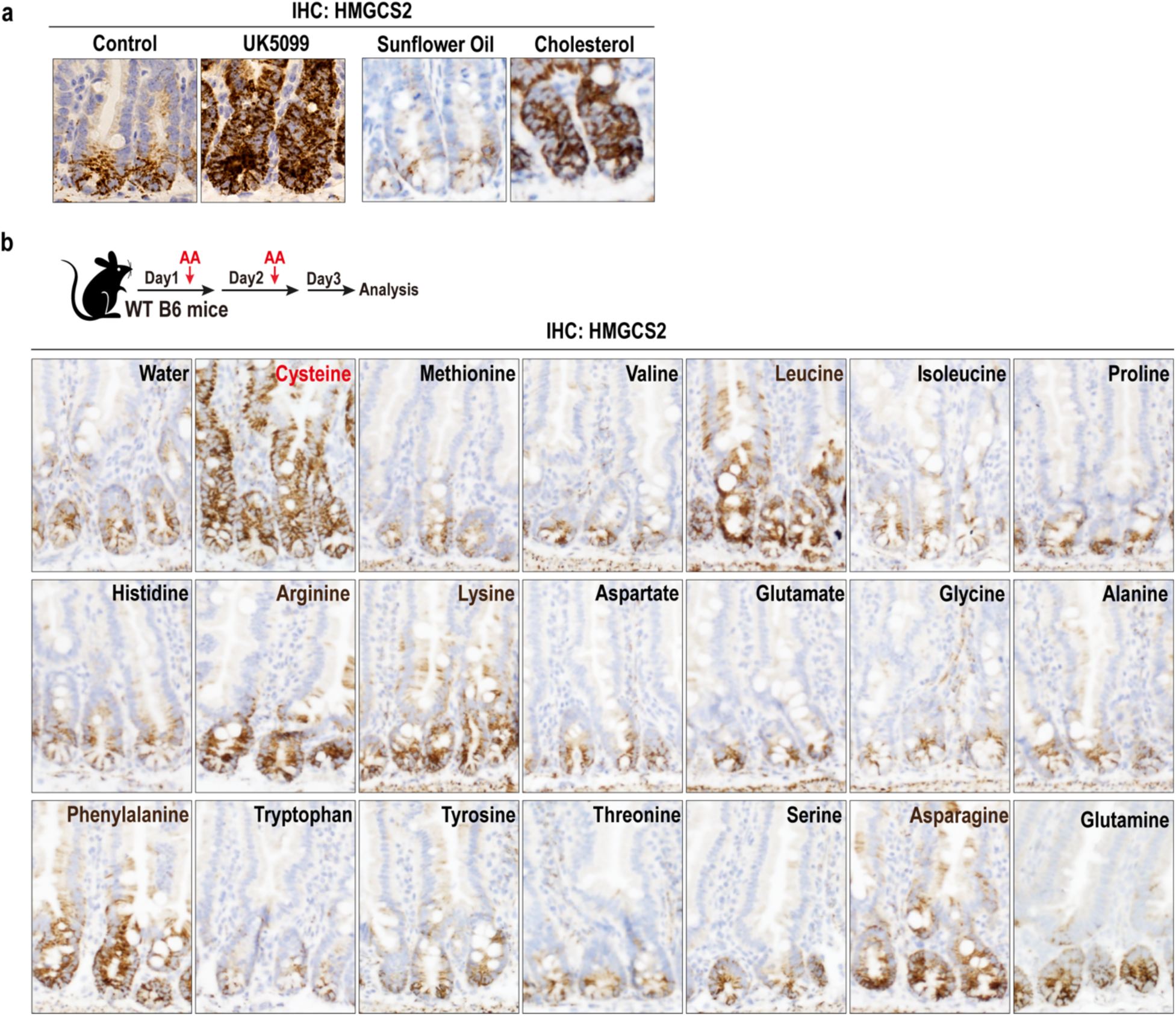
Amino acids show various impacts on ISC function. **a**, HMGCS2 IHC in small intestinal crypts from the mice treated with mitochondrial pyruvate carrier inhibitor UK5099^4^ and cholesterol^6^. **b**, Schematic (top) of each amino acid treatment, and HMGCS2 IHC (bottom) in small intestinal crypts from the mice 3 days after treated with each of 20 proteinogenic amino acids *in vivo*.

### Cysteine boosts ISC-mediated regeneration

We next sought to test how cysteine affects ISC *in vitro* and *in vivo* function by developing a cysteine-rich diet (CysRD) with 40 grams of cystine (the stable oxidized derivative of two cysteine molecules connected by a disulfide bond) per kilogram and a calorie-matched control diet with 3 grams of cystine (**Extended Data Fig. 2a**). Cystine is the predominant form of extracellular cysteine and accounts for the majority of how cysteine is acquired by cells^27^. Mice fed a CysRD for 6 weeks appeared healthy and had no effects on blood glucose or ketone levels (**Extended Data Fig. 2b**). Although both cohorts exhibited similar food uptake, mice fed a CysRD had slower bodyweight gain and less visceral fat accumulation (**Extended Data Fig. 2c-d**). The CysRD had no impact on intestinal proliferation *in vivo* as assessed by BrdU incorporation, villi length, crypt depth, and ISC numbers based on OLFM4^+^ IHC (an alternate Lgr5^+^ ISC marker) and Lgr5-eGFP (using *Lgr5-EGFP-IRES-Cre^ERT2^* mouse model^28^) flow cytometry quantification (**Extended Data Fig. 3a-e**). However, small intestinal crypts isolated from CysRD-fed mice or mice orally administered cysteine for 3 days had markedly enhanced organoid-forming efficiency compared to control mice, demonstrating that cysteine supplementation primes intestinal stem cell activity in conditions that mimic regeneration (i.e., the ability of ISCs and early progenitors to engender organoid formation in culture) (**Fig. 2a-c, Extended Data Fig. 3f-g**).

**Fig. 2.**
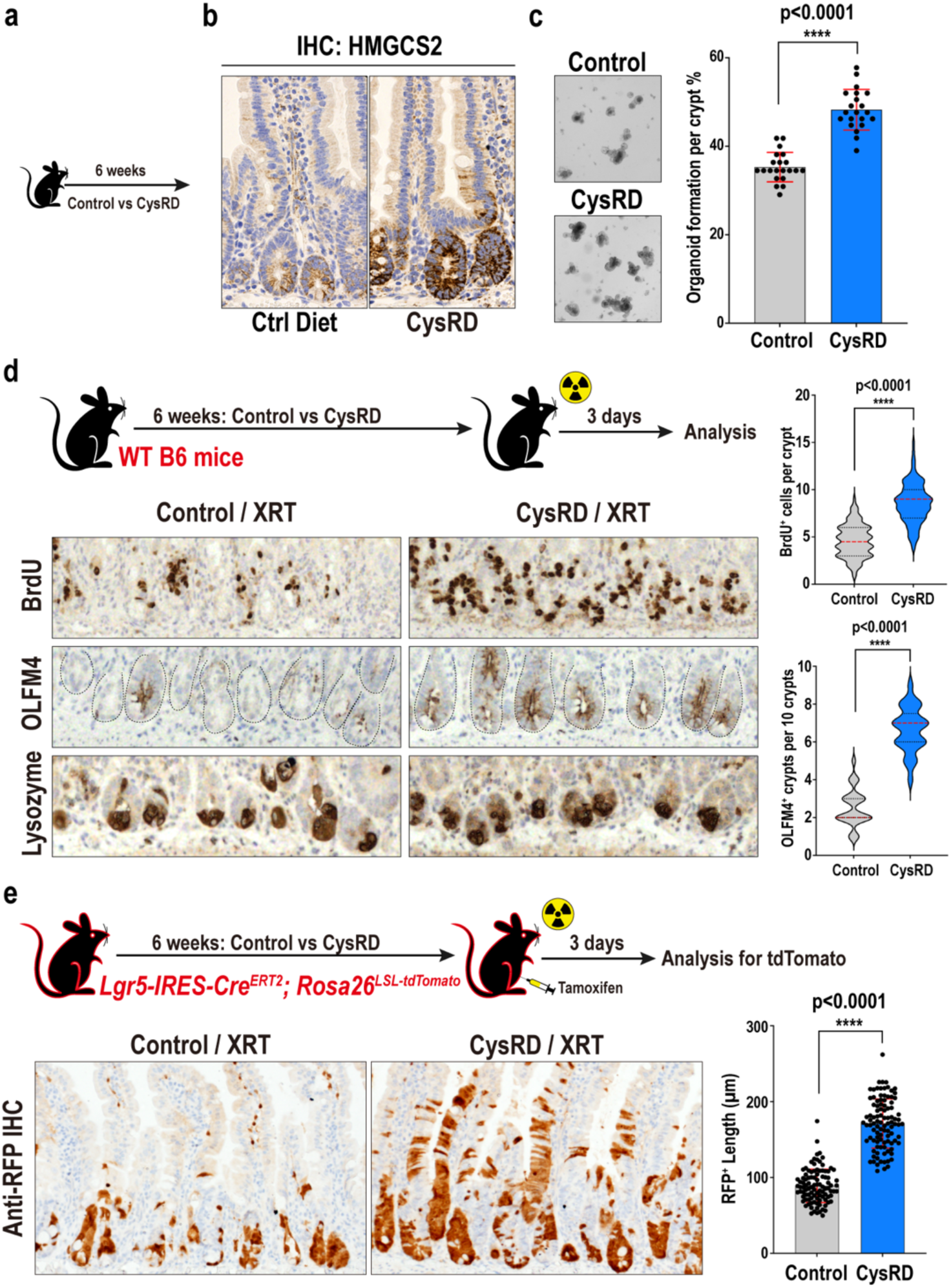
Cysteine boosts ISCs-mediated regeneration. **a**, Schematic of cysteine-rich diet (CysRD) dietary regimen in WT B6 mice. **b**, IHC for HMGCS2 in the intestinal crypts from control and CysRD-fed mice. **c**, Representative images and quantification of day-3 organoids formed by small intestinal crypts from control and CysRD-fed mice. n = 21 biological replicates per group. **d**, Schematic (top) of the intestinal regeneration assay, including the timeline of dietary regimen, irradiation (XRT, 7.5Gy x 2), and tissue collection. Representative images (bottom) of regenerating crypts by IHC for 4 hours pulse of BrdU cell proliferation and OLFM4 stem cell markers. Lysozyme labels post-mitotic Paneth cells. Quantification (right) of day-3 BrdU^+^ cells (per crypt) and OLFM4^+^ crypts (per 10 crypts) from 5 control and 5 CysRD diets treated mice intestine. 20 data points for BrdU^+^ cells and 5 data points for OLFM4^+^ crypts per mouse were collected. **e**, Schematic (top) of the intestinal regeneration assay including the timeline of dietary regimens, irradiation (XRT, 7.5Gy x 2), and tissue collection, in the Lgr5 lineage tracing model with *Lgr5-IRES-Cre^ERT2^; Rosa26^LSL-tdTomato^* reporter mice. Representative images (bottom) of IHC for tdTomato^+^ Lgr5^+^ ISC-derived progenies in the small intestine. Quantification (right) of day-3 RFP^+^ length from 5 control and 5 CysRD diets treated mice intestine. 20 data points per mouse were collected. Unpaired two-tailed Student’s t-tests (**c**,**d**,**e**). Data are mean ± s.d.

To ascertain whether CysRD also boosted ISC function *in vivo*, we damaged the intestine with lethal irradiation and enumerated numbers of regenerative crypts 3 days post-irradiation. CysRD-fed mice had higher numbers of proliferating BrdU^+^ ISCs and progenitors than controls, while Lysozyme^+^ Paneth cell numbers were unaffected (**Fig. 2d**). We then directly assessed how the same injury model directly impacted *in vivo* ISC output using the Lgr5 ISC lineage-tracing (*Lgr5-IRES-Cre^ERT2^*; *Rosa26^LSL-tdTomato^*) mouse model^29^. In this model, tamoxifen administration activates tdTomato in Lgr5^+^ ISCs, thereby indelibly labeling ISCs and their progeny. Using this model, we labeled ISCs 24 hours before radiation exposure and then visualized tdTomato-positive cells 3 days after irradiation (**Fig. 2e**). CysRD feeding dramatically supported ISC-mediated repair after radiation-induced injury, as revealed by significantly increased numbers of tdTomato^+^ cells, indicating that, in response to injury, cysteine treatment augments ISC proliferation and output (**Fig. 2d-e**). These data, thus, illustrate that cysteine boosts the regenerative capacity of ISCs.

### Cysteine mediates its ISC-enhancing effects through CD8αβ^+^ T cells

We then queried how cysteine actuates its effects on ISCs. Although small intestinal histology was unremarkable, we noticed increased crypt CD8^+^ T cell intraepithelial lymphocytes (IELs) (**Fig. 3a-b**), demonstrating that dietary cysteine alters the ISC immune microenvironment. We employed flow cytometry and immunofluorescence to further characterize these IELs under homeostatic and irradiation-induced injury conditions. We observed a 2- to 3-fold increase in CD8αβ^+^ T cell numbers within the small intestinal crypts and lamina propria, but not in the colon, of CysRD-fed mice compared to controls (**Fig. 3c**, **Extended Data Fig. 4a**). Other immune cell populations, such as CD4^+^ T cells, CD8αα^+^ T cells, CD19^+^ B cells, and NK1.1^+^ NK cells, were unaltered with CysRD feeding (**Fig. 3c**). CysRD feeding, interestingly, augments CD8αβ^+^ T cell numbers in homeostasis and repair after damage (**Fig. 3d**). We then fed Rag2^−/−^ (KO) mice^32^, which lack mature T and B cells, with control or CysRD for 6 weeks and then damaged the intestines with irradiation. Notably, the absence of T and B cells nullified cysteine’s regenerative effects on ISCs (**Fig. 3e**). Given that CysRD feeding boosted intraepithelial CD8αb^+^ T cell numbers, we investigated whether intestinal immune cells are essential for mediating cysteine’s regenerative effects on ISCs. Intraepithelial CD8αβ^+^ T cells from control and CysRD crypts were co-cultured with untreated Lgr5-GFP ISCs in the *in vitro* organoid assay. While control CD8αβ T cells slightly enhanced the organoid-forming capacity of ISCs, CD8αβ T cells from the CysRD group had a dramatic effect on ISC organoid-forming capacity (**Fig. 3f**) signifying that dietary cysteine mediates its ISC-enhancing effects through CD8αβ^+^ T cells.

**Fig. 3.**
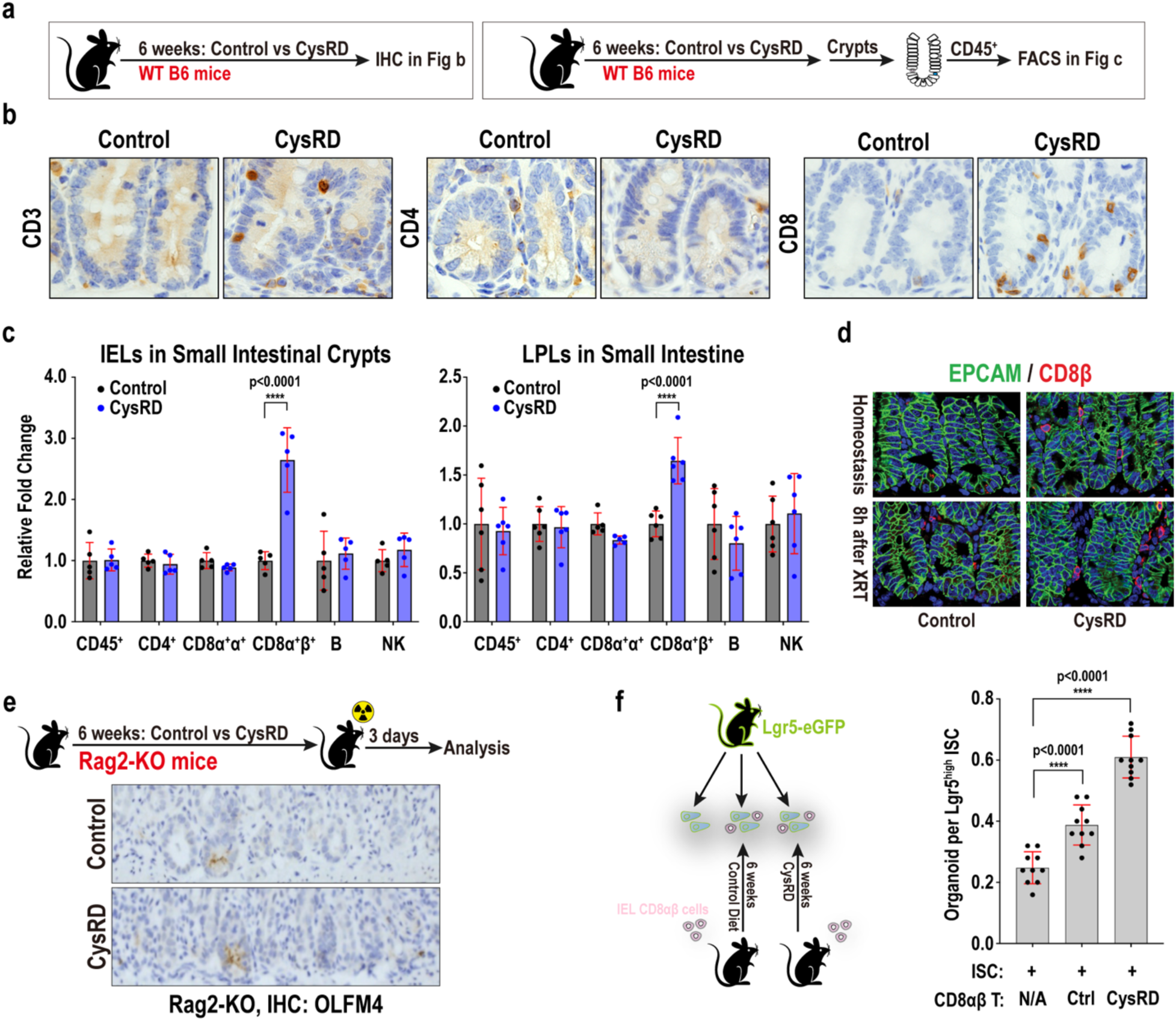
Cysteine expands crypt-associated CD8αβ T cells and their ISC-enhancing ability. **a**, Schematic of mouse dietary regimen, tissue processing, and analysis. **b**, IHC for CD3, CD4, and CD8 in the small intestinal crypts from control and CysRD-fed mice. **c**, Relative fold changes of small intestine intraepithelial (IEL) and lamina propria lymphocytes (LPLs) from control and CysRD-fed mice. **d**, Representative crypt images of EPCAM (epithelial cells) and CD8αβ cells from control and CysRD fed mice before and 8 hours after irradiation. **e**, Schematic (top) of the intestinal regeneration assay, including the timeline of dietary regimen, irradiation (XRT, 7.5Gy x 2), and tissue collection in the Rag2^−/−^ mice. IHC for regenerating crypts using the OLFM4 stem cell marker after irradiation mediated tissue damage and repair in Rag2^−/−^ mice from the control and CysRD-fed mice. **f**, Schematic (left) and quantification (right) of organoid formation assay using isolated Lgr5-GFP^high^ ISCs with intestine intraepithelial CD8αβ T cells from control and CysRD fed mice. 10ul organoid culture droplet contains 2500 Lgr5-GFP^high^ ISCs and 2000 CD8αβ^+^ T cells. Unpaired two-tailed Student’s t-tests (**c, f**). Data are mean ± s.d.

### CD8αβ^+^ T cell-derived IL-22 boosts ISC-mediated repair

To investigate the role of cytokines in mediating the effects of cysteine on ISC-driven tissue repair, we conducted an unbiased cytokine array assay, profiling 115 mouse cytokines in the small intestine following CysRD feeding. Among the top candidates, interleukin-22 (IL-22) emerged, displaying a significant elevation after 6 weeks of CysRD treatment (**Fig. 4a**). IL-22, a cytokine known to promote ISC-mediated intestinal epithelial regeneration, is predominantly produced by lymphocyte populations, including innate lymphoid cells type 3 (ILC3), CD4^+^ T cells, CD8^+^ T cells, γδ T cells, and natural killer (NK) cells^20,21,33,34,35^. Interestingly, isolated crypts derived from CysRD-fed mice exhibited a 3-fold increase in IL-22 levels. Notably, crypts with depleted CD8αβ^+^ T cells showed IL-22 levels similar to control diet (**Fig. 4b**). Furthermore, IL-22 production was significantly upregulated in CD8αβ^+^ T cells following CysRD feeding (**Fig. 4c**). A single intraperitoneal (i.p.) injection of IL-22 was sufficient to induce HMGCS2 expression in crypt cells (**Fig. 4d**), highlighting a mechanism whereby cysteine induces IL-22 production in CD8αβ^+^ T cells, which in turn activates HMGCS2 expression in ISCs and progenitor cells.

**Fig. 4.**
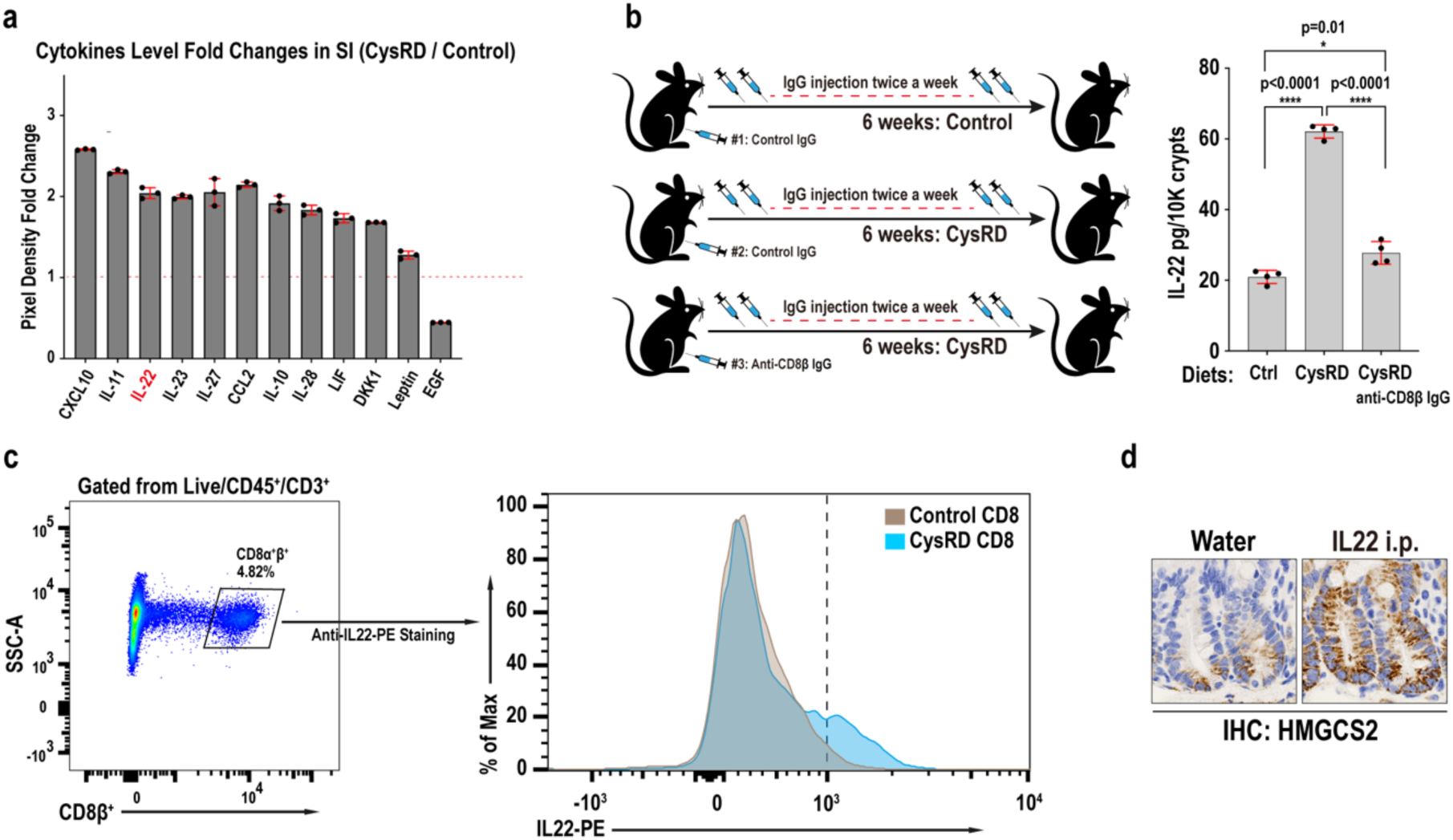
CD8αβ^+^ T cell-derived IL-22 mediates cysteine’s pro-regenerative effects. **a**, Relative fold changes of representative highly expressed cytokines that respond to CysRD feeding (6 weeks) in the small intestine tissue using mouse XL cytokine array. **b**, Schematic (left) and quantification (right) of intestinal crypts IL-22 levels from control and CysRD fed mice using IL-22 ELISA assay. **c**, Flow cytometry for intestinal intraepithelial CD8αβ^+^ T cells gating panel (left), and anti-IL-22-PE histogram (right) in control and CysRD fed mice. **d**, Small intestinal crypts HMGCS2 IHC from control and IL-22 i.p. injected mice. Unpaired two-tailed Student’s t-tests (**b**). Data are mean ± s.d.

## Discussion

Here, we describe how dietary cysteine augments intestinal stemness non-autonomously via CD8αβ^+^ T cells, an immune niche population not previously implicated in coordinating micronutrients with ISC activity. A CysRD diet treatment promotes the expansion of crypt-associated CD8αβ^+^ T cells (**Fig. 3**). This, in turn, boosts their production of IL-22, a trophic factor that augments Lgr5^+^ ISCs’ ability to repair intestinal injury^19,20,21^ (**Fig. 4**). Amino acids serve as essential building blocks for proteins and nucleotides and as regulatory molecules in cell signaling pathways and precursors for bioactive ligands and cofactors^36^. Previous studies have shown that a deficiency in certain amino acids can significantly impact stem cell behavior^37,38,39,40,41,42,43,44^. However, the supplementation of specific amino acids and their effects on stem cells and tissue regeneration are less understood. Recent research has proposed that cysteine is a critical regulator of mitochondrial function, fasting-induced tissue responses, and body fat metabolism through diverse mechanisms^45,46,47^. Our study uncovers a previously unrecognized mechanism whereby dietary cysteine stimulates ISC function in injury by coupling cysteine metabolism between epithelial and CD8αβ^+^ T cells.

Given that we find that dietary cysteine enhances ISC activity in injury, our findings raise the possibility that cysteine supplementation may benefit conditions for which a healthier ISC and progenitor pool improves epithelial barrier integrity and regeneration. For example, cysteine supplementation may improve ISC repair of the epithelial barrier in patients with acute enteritis from infectious etiologies, radiation exposure, or cytotoxic chemotherapeutics that damage the small intestinal epithelium. It will also be interesting to understand why cysteine improves small intestinal regeneration but not the colon (**Extended Data Fig. 4**) and whether cysteine supplementation has analogous effects on other tissue stem cells. Finally, although cysteine and its antioxidant derivatives, such as glutathione and taurine, are known for their anti-aging properties^48,49^, cysteine supplementation via the mechanism described here may ameliorate dampened regeneration in the elderly intestine where both ISC and CD8αβ^+^ T cell functions are reduced.

## Acknowledgements

We thank the Swanson Biotechnology Center at the Koch Institute, including the Flow Cytometry, Histology, and ES Cell and Transgenics core facilities; the Department of Comparative Medicine for mouse husbandry support; S. Holder and members of the Hope Babette Tang (1983) Histology Facility for substantial histological support; all the members of the Yilmaz laboratory for discussions; K. Kelley for laboratory management; and L. Galoyan for administrative assistance. Ö.H.Y. is supported by the National Institutes of Health (R01CA211184, R01CA034992, R01CA257523, R01DK126545, U01CA250554 and U54CA224068); a Pew-Stewart Trust scholar award; the Kathy and Curt Marble cancer research award; a Koch Institute–Dana-Farber/Harvard Cancer Center Bridge project grant; and AFAR. Ö.H.Y. receives support from the MIT Stem Cell Initiative. F.C. is supported by a Damon Runyon Postdoctoral Fellowship Award (DRG-2463-22). Q.Z. is supported by a Cancer Research Institute Postdoctoral Fellowship (CRI4478). J.E.S. is supported by National Institutes of Health F32 and P30 Fellowships (F32DK128872, P30-DK040561). S.S. is supported by Crohn’s & Colitis Foundation Award (CCFA-623914) and American Heart Association Award (19POST34380588). Y.M.S. is supported by National Institutes of Health (R01CA245546).

## Author Contributions

F.C. initiated the project, conceived, designed, performed, and interpreted all the experiments and wrote the manuscript with help from Ö.H.Y.. Q.Z., J.E.S., Y.Y., Z.Y., H.S. performed experiments and assisted with data collection. J.T.H.S. assisted with the intestinal tissue metabolomic analysis. S.S., Y.M.S. provided research materials. J.A. assisted with immune cell experimental design and data interpretation. All the authors assisted in interpreting the experiments, writing, and editing the paper.

## Ethics declarations

Competing interests: The authors declare no competing interests.

## Methods

### Mice

Mice were under the husbandry care of the Department of Comparative Medicine in the Koch Institute for Integrative Cancer Research. All procedures were conducted in accordance with the American Association for Accreditation of Laboratory Animal Care and approved by MIT’s Committee on Animal Care. The following strains were obtained from the Jackson Laboratory: *Lgr5-eGFP-IRES-CreERT2* (strain name: B6.120P2-Lgr5tm1(cre/ERT2)Cle/J, stock number 008875), Rag2^−/−^ (strain name: B6.Cg-*Rag2^tm1.1Cgn^*/J, stock number 008449). The *Lgr5-IRES-Cre^ERT2^*; *Rosa26^LSL-tdTomato^* mice were a gift from H. Clevers and has been previously described^7,29^. High cysteine diet (Research Diets D21081608) was provided to male and female mice at the age of 8-12 weeks for 6 to 8 weeks. Control mice were provided a purified control diet (Research Diets D12450J). In Figure 1, the mice were on standard chow at the time of amino acid treatment, and each amino acid was given to the mice with oral gavage at the following doses: Arginine 740mg/Kg, Cysteine 80mg/Kg, Histidine 160mg/Kg, Isoleucine 260mg/Kg, Leucine 150mg/Kg, Lysine 460mg/Kg, Methionine 80mg/Kg, Phenylalanine 170mg/Kg, Threonine 270mg/Kg, Tryptophan 50mg/Kg, Tyrosine 240mg/Kg, Valine 260mg/Kg, Glycine 90mg/Kg, Alanine 50mg/Kg, Asparagine 50mg/Kg, Aspartate 50mg/Kg, Glutamate 50mg/Kg, Proline 100mg/Kg, Serine 130mg/Kg, Glutamine 150mg/Kg, Taurine 200mg/Kg, GSH (i.p. & oral gavage) 200mg/Kg, NAC (i.p. & oral gavage) 200mg/Kg. All mice were sex and aged matched and provided food *ad libitum*. Floxed alleles were excised following intraperitoneal injection of tamoxifen suspended in 1 part 100% ethanol: 9 parts sunflower seed oil (Spectrum S1929) at a concentration of 10 mg/ml and dosed at 100mg/kg. Mice harboring conditional alleles were administered tamoxifen two times within one week unless otherwise specified.

### Irradiation experiments

Mice were challenged by a lethal split dose of irradiation using gamma-cell irradiator using Cs as irradiation source, 7.5Gy x 2 with six hours between exposures. Tissue was collected 72 hours post the last dose. Numbers of surviving crypts were enumerated in the jejunum from OLFM4 stained tissue.

### *In vivo* CD8β^+^ T cells depletion

To deplete CD8β^+^ T cells, 200 μg of control IgG (BioXcell, BE0088) or anti-CD8β (BioXcell, BE0223) in dilution buffer (BioXCell, IP0070) was intraperitoneally injected twice a week respectively. Antibody injection was initiated 2 days before the CysRD treatment and sustained during the entire dietary intervention.

### Crypt isolation and culture

Small intestines were removed, flushed with PBS^−/−^ (no calcium, no magnesium), opened laterally, gently wiped to remove mucus layer and cut into ∼1cm sections. Intestine pieces were rinsed 3 times and incubated at 4°C in PBS^−/−^ with 10 mM EDTA for 45 min. Crypts were then mechanically separated from the connective tissue by shaking and filtered through 70μm mesh into a 50 mL conical tube to remove villi and tissue fragments. Isolated crypts were counted and embedded in 3.5:6.5 media: Matrigel (Corning 356231 growth factor reduced) mixture at 5-10 crypts per μl and cultured in a crypt culture medium. Unless otherwise described, crypts were grown in Advanced DMEM (GIBCO, 12491015) supplemented with EGF 40 ng/ml (Peprotech, 315-09), Noggin 200 ng/ml (Peprotech 250-38), R-Spondin 500 ng/ml (Peprotech, 315-32), N-acetyl-L-cysteine 1 μM (Sigma Aldrich, A9165), B27 1x (Life Technologies, 17504044), Chir99021 1 μM (LC Laboratories, C-6556), Y-27632 dihydrochloride monohydrate 10 μM (Sigma Aldrich, Y0503). Intestinal crypts were plated in 10 (5 μl) droplets of Matrigel and placed on flat bottomed 48 (96) well plates (Olympus 25-108 (48) 25-109 (96)) and allowed to solidify for 15 min in a 37°C incubator. 250/150 μL of crypt medium was added to each well and maintained at 37°C in a humidified incubator at 5% CO2. Crypt medium was changed every three days. Clonogenicity (colony-forming assay) was assessed on day 3. Organoid images were acquired on Nikon Ti Eclipse epifluorescence microscope equipped with LCD light source (lumencor light engine® SOLA, SM 5-LCR-SA) and sCMOS camera (Andor Zyla 4.2 cMOS). For the co-culture assay, flow-isolated Lgr5^hi^ ISCs and CD8αβ^+^ T cells cells were centrifuged at 200g for 3 min, resuspended, mixed (2500 ISCs + 2000 CD8αβ^+^ T) in the 3 µl of crypt medium and seeded onto 7 µl Matrigel containing 1 µM JAG-1 protein (AS-61298, AnaSpec) in a flat bottom 48-well plate (3548, Corning). The Matrigel and cells mixture were allowed to solidify before adding 300 µl of crypt culture medium.

### Immunohistochemistry (IHC) and immunofluorescence (IF)

Tissues were fixed in 10% formalin, paraffin embedded and sectioned in 4-5micron sections as previously described (Mana et al, 2021). Antigen retrieval was performed using Borg Decloaker RTU solution (Biocare Medical, BD1000G1) and a pressurized Decloaking Chamber (Biocare Medical, NxGen). Antibodies and respective dilutions used for immunohistochemistry are as follows: rabbit monoclonal anti-OLFM4 (1:10,000, CST, 39141), rat anti-BrdU (1:2000, Abcam 6326), rabbit polyclonal anti-lysozyme (1:2000, Thermo RB-372-A1), and rabbit monoclonal anti-HMGCS2 (1:1000, Abcam ab137043). Biotin-conjugated secondary donkey anti-rabbit, or anti-rat antibodies were used (1:500, Jackson ImmunoResearch). Vectastain Elite ABC immunoperoxidase detection kit (Vector Laboratories, PK6100) was followed by Signalstain® DAB substrate kit for visualization (CST 8049S). All antibody dilutions were performed in Signalstain® Antibody Diluent (CST 8112L). The following primary antibodies were used for immunofluorescence in snap frozen tissue section: rabbit polyclonal anti-EPCAM (1:500, Abcam ab71916), rat monoclonal anti-CD8β (1:1000, BioXCell BE0223). Alexa Fluor secondary antibodies, anti-rabbit 488, and anti-rat 568, were used for visualization. Tissue was mounted using Invitrogen Prolong Gold mounting medium containing DAPI. Images were acquired using a 60x objective using an Olympus FV1200 Laser Scanning Confocal Microscope.

### Immunoblotting

The following antibodies were used: rabbit monoclonal anti-HMGCS2 (1:1000, Abcam ab137043), rabbit monoclonal anti-phospho-S6 Ribosomal Protein (Ser235/236) (1:1000, CST 4858), monoclonal rabbit anti-S6 Ribosomal Protein (1:1000, CST 2217) and mouse monoclonal anti-Actin (1:10000, Sigma MAB1501). 20ug of crypt lysates were loaded per sample onto a 4%–12% gradient gel, transferred on to PVDF membrane (Immobilon-P transfer, Millipore, ipvh00010), and analyzed using IgG-HRP antibodies (1:3000, CST, 7076, 7074) and Advansta WesternBright ECL detection kit (K-12045-D20).

### Flow cytometry

The isolated crypts were dissociated to individual cells with TrypLE Express (Invitrogen), 32 °C for 1 min. For ISC isolation, an antibody cocktail comprising CD45-PE (eBioscience, 30-F11), CD31-PE (Biolegend, Mec13.3), Ter119-PE (Biolegend, Ter119), and EPCAM-APC (eBioscience, G8.8) was used for cell staining. For crypt associated immune cell isolation, an antibody cocktail comprising (1) CD45-APC-Cy7 (BioLegend, 30-F11), CD3-FITC (BioLegend, 17A2), CD8b-APC (BioLegend, YTS156.7.7), CD19-BUV737 (BD Biosciences, 1D3), NK1.1-Alexa Flour 700 (BD Biosciences, PK136); (2) CD3-FITC (BioLegend, 17A2), CD8a-PE (BioLegend, QA17A07), CD8b-APC (BioLegend, YTS156.7.7); (3) EPCAM-PE (eBioscience, G8.8), CD8b-APC (BioLegend, YTS156.7.7) were used. The antibody staining process was at 4 °C for 20 min in dark. The cells were washed and resuspended in FACS buffer (RPMI 1640 with 2% FBS) with DAPI (Thermo FScientific, 62248; 1:10,000 dilution) or 7AAD (Invitrogen, A1310; 1:500 dilution). For intracellular staining, cells were fixed for 30 min on ice in BD Cytofix/Cytoperm Fixation/Permeabilization (BD Biosciences, 554722) and washed in BD Perm/Wash (BD Biosciences, 554723) diluted 1:10 in ddH_2_O. Intracellular staining of IL-22-PE (BioLegend, Poly5164) was performed in Perm/Wash buffer for 30 min at 4 °C. Cells were washed and resuspended in Perm/Wash buffer.

To isolate immune cells from lamina propria, the small intestines were dissected, flushed with PBS^−/−^ (no calcium, no magnesium), opened laterally, gently wiped to remove mucus layer and cut out the Peyer patches. The intestines were cut into ∼1cm sections, incubated in PBS-EDTA (10mM) at 37°C for 30 min. The intestine epithelial cells were removed out by vigorously shaking for 3 times in the PBS^−/−^. The remaining tissues were minced into 1-2mm pieces, incubated with 20ml pre-warmed 2% FBS RPMI medium containing 20ug/ml Liberase DH and 50ug/ml DNase I at 37°C, on a ThermoMixer at 1000rpm for 30min. 15ml supernatant were collected and stored on ice. For the remaining 5ml, repeatedly draw the mixture through an 18-gauge needle using a syringe to further mince the intestinal pieces. Add 15ml pre-warmed 2% FBS RPMI medium containing 20ug/ml Liberase DH and 50ug/ml DNase I at 37°C, on a ThermoMixer at 1000rpm for 30min. The 35ml supernatant were combined and filtered through a 70um strainer. The filtered samples were centrifuged at 4°C, 500g for 5min, and the pellets were resuspended with 10ml 5% FBS HBSS, filtered through 45um into new tubes for downstream FACS analysis. The aforementioned antibody cocktail comprising CD45-APC-Cy7 (BioLegend, 30-F11), CD3-FITC (BioLegend, 17A2), CD8b-APC (BioLegend, YTS156.7.7), CD19-BUV737 (BD Biosciences, 1D3), NK1.1-Alexa Flour 700 (BD Biosciences, PK136) were then used to detect immune cell changes in mice fed with control and high-cysteine diets.

### Metabolomics analysis

The small intestine tissues were quickly dissected out, washed twice with 150mM Ammonium acetate, and snap-freeze into liquid nitrogen. The snap-freezed tissues were homogenized into powder using mortar-pestle in liquid nitrogen. About 20mg of the tissue powder was used for metabolites extraction in 80% MeOH with OMNI bead homogenizer. The tissues/80%MeOH mixture was incubated for 24 hours at −80°C, spined at the top speed (16,000 g) for 20 minutes at 4°C. The MeOH supernatants were transferred into new tubes and placed on ice. The sample pellets were resuspended in 500μl 0.2M NaOH, heated at 95°C for 10 minutes, and the protein concentration was measured by BCA assay using 1:10 dilution. Finally, the 500 μg protein equivalent of MeOH supernatants were dried using multivap nitrogen evaporator.

Dried metabolites were resuspended in 50% ACN:water at 50 mg tissue extract/ml. 5μl was loaded onto a Luna NH2 3um 100A (150 × 2.0 mm) column (Phenomenex) using a Vanquish Flex UPLC (Thermo Scientific). The chromatographic separation was performed with mobile phases A (5 mM NH4AcO pH 9.9) and B (ACN) at a flow rate of 200 μl/min. A linear gradient from 15% A to 95% A over 18 min was followed by 7 min isocratic flow at 95% A and reequilibration to 15% A. Metabolites were detected with a Thermo Scientific Q Exactive mass spectrometer run with polarity switching in full scan mode using a range of 70-975 m/z and 70.000 resolution. Maven (v 8.1.27.11) was used to quantify the targeted polar metabolites by AreaTop, using expected retention time and accurate mass measurements (< 5 ppm) for identification. Data analysis, including principal component analysis and heat map generation was performed using in-house R scripts.

### Statistics and reproducibility

Unless otherwise specified in the main text or figure legends, all experiments reported in this study were repeated at least three independent times. Unless otherwise specified in the main text or figure legend, all sample numbers (n) represent biological replicates. For the organoid assays, 3–5 wells per group with at least 3 different mice were analysed. No sample or animals were excluded from analysis, and sample size estimates were not used. Studies were not conducted blind with the exception of all histological analyses. Age-matched and sex-matched mice were randomly assigned to groups. All values are presented as mean ± s.d. Unless otherwise specified in the figure legend, intergroup comparison was performed using unpaired two-tailed t-tests. Statistical analysis was performed using GraphPad Prism 9. Statistical details can be found in the figure legends.

---

**Extended Data Fig. 1.**
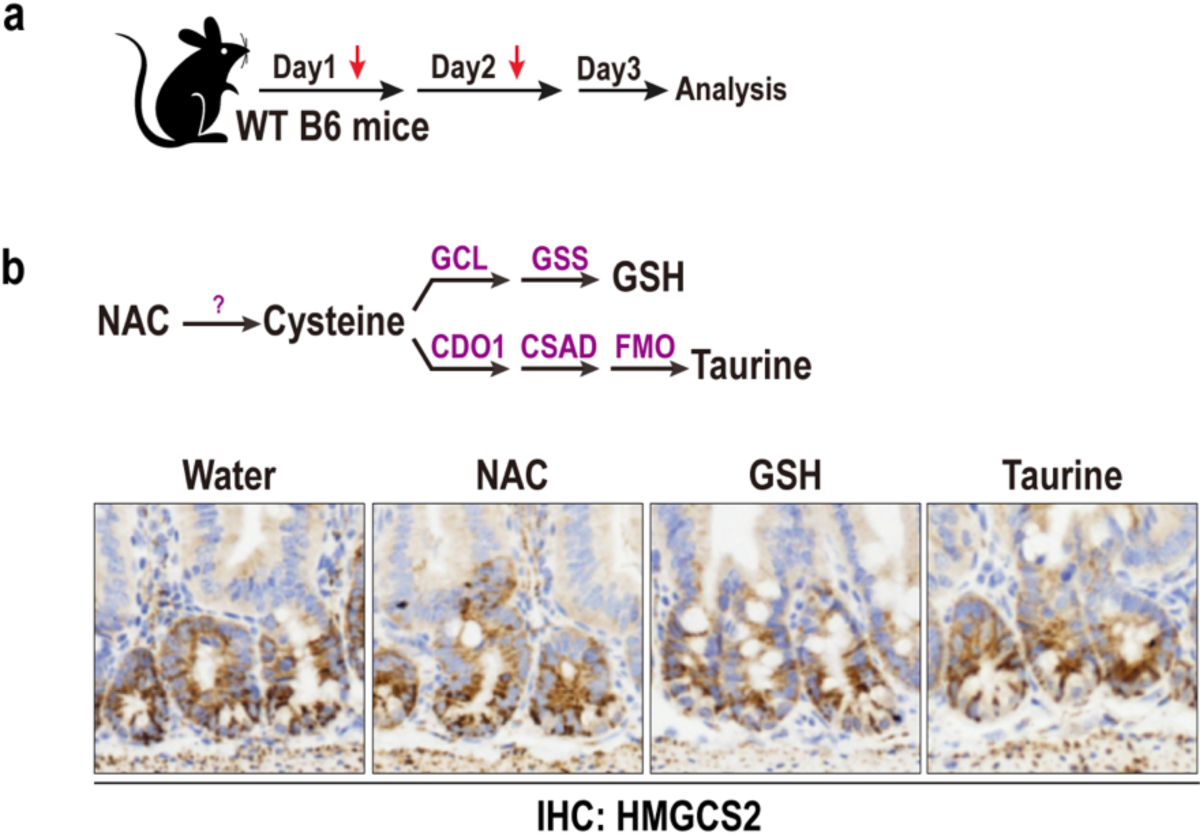
Effect of antioxidant derivatives of cysteine metabolism on HMGCS2 expression. **a**, Schematic of nutrients treatment *in vivo*. **b**, N-acetyl cysteine (NAC), Glutathione (GSH), and taurine are well-established metabolites associated with cysteine metabolism. HMGCS2 IHC in small intestinal crypts treated with NAC, GSH, and taurine by i.p. and oral gavage.

**Extended Data Fig. 2.**
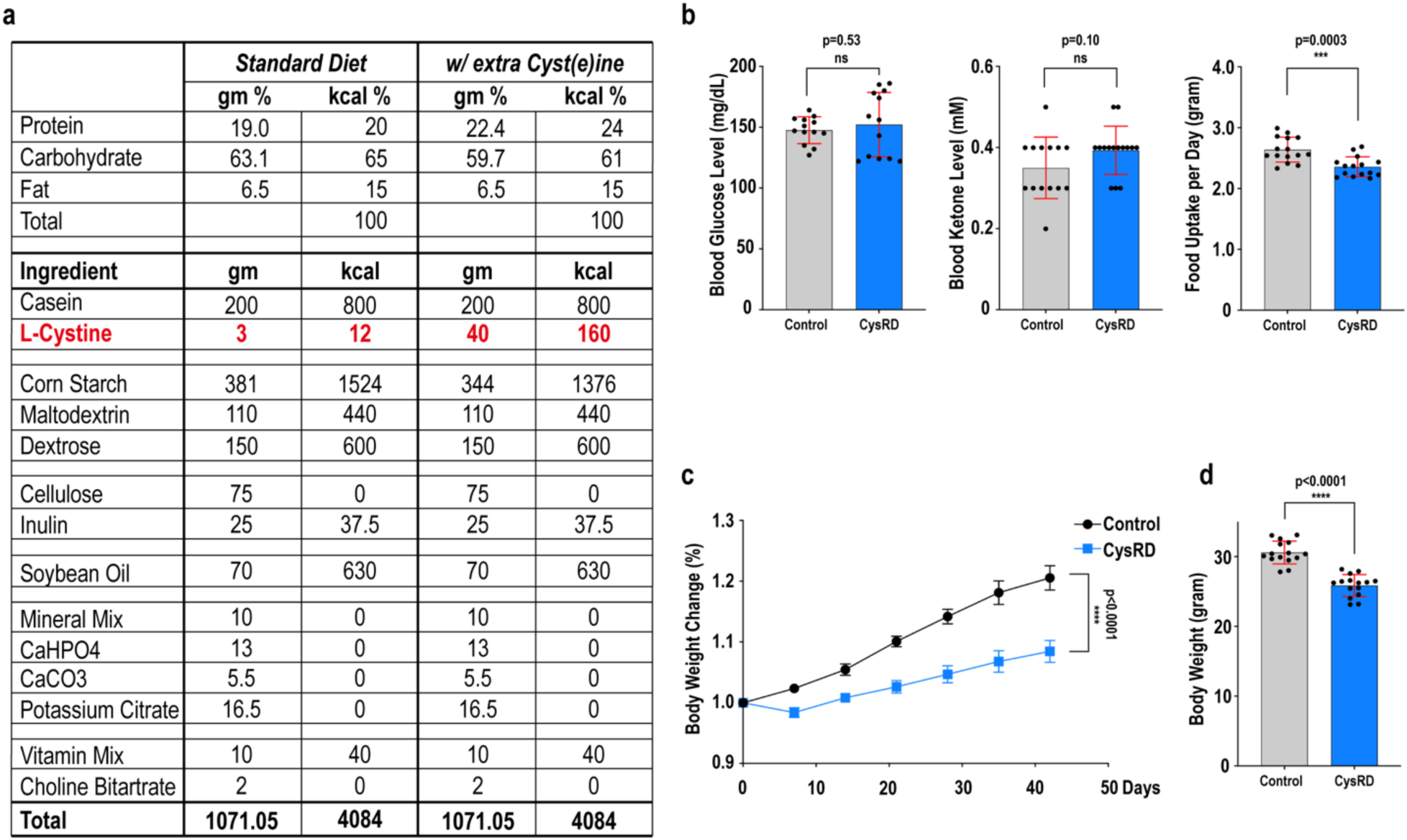
Cysteine-rich diet composition and mouse physiology. **a**, Control and cysteine-rich diet (CysRD) composition. **b**, Mouse blood glucose and ketone levels after 6 weeks of control and CysRD feeding. Mouse daily food uptake during the control and CysRD feeding. **c-d**, Relative body mass changes (**c**) and terminal body mass (**d**) for mice fed with 6 weeks of control diet and CysRD. Unpaired two-tailed Student’s t-tests (**b**,**c**,**d**). Data are mean ± s.d.

**Extended Data Fig. 3.**
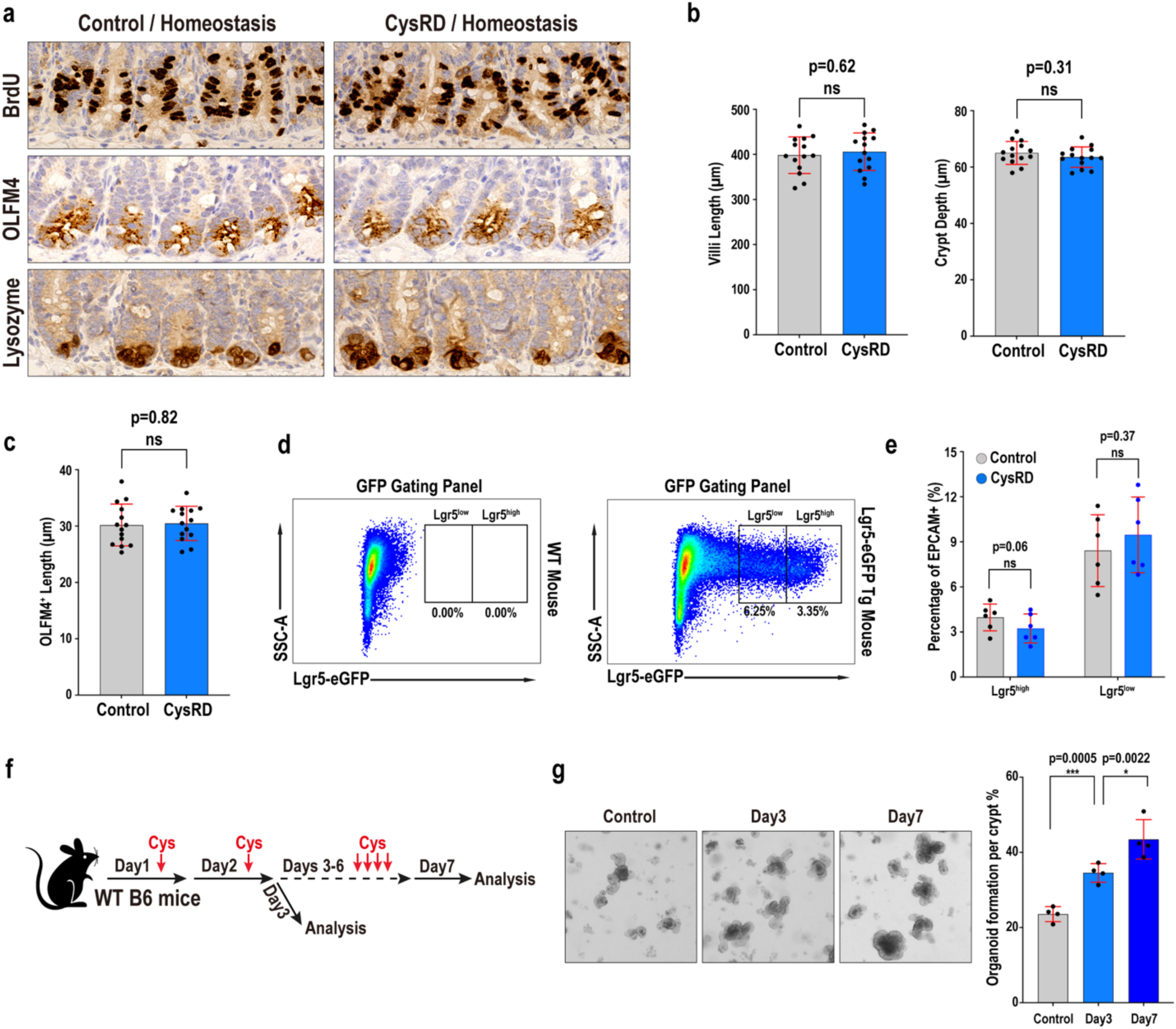
Effects of Cysteine-rich diet on ISC and progenitor cells. **a**, IHC detection for intestinal stem cell and progenitor cell proliferation (4-hour BrdU pulse), stem and progenitor cell numbers (OLFM4), and post-mitotic Paneth cell identity (Lysozyme) from control diet and CysRD fed mice in homeostasis. **b**, Villi length and crypt depth in the small intestine isolated from control-diet and CysRD-fed mice in homeostasis. **d-e**, Flow cytometry for Lgr5-GFP gating panel (**d**) and quantification (**e**) of ISCs (Lgr5-GFP^hi^) and progenitors (Lgr5-GFP^low^) frequency from control diet and CysRD 6 week fed mice. n = 6 for each group. **f-g**, Schematic of the short-term cysteine treatment by oral gavage (**f**), followed by *in vitro* crypt organoid formation assay (**g**). n = 4 biological replicates per group. Unpaired two-tailed Student’s t-tests (**b**,**c**,**e,g**). Data are mean ± s.d.

**Extended Data Fig. 4.**
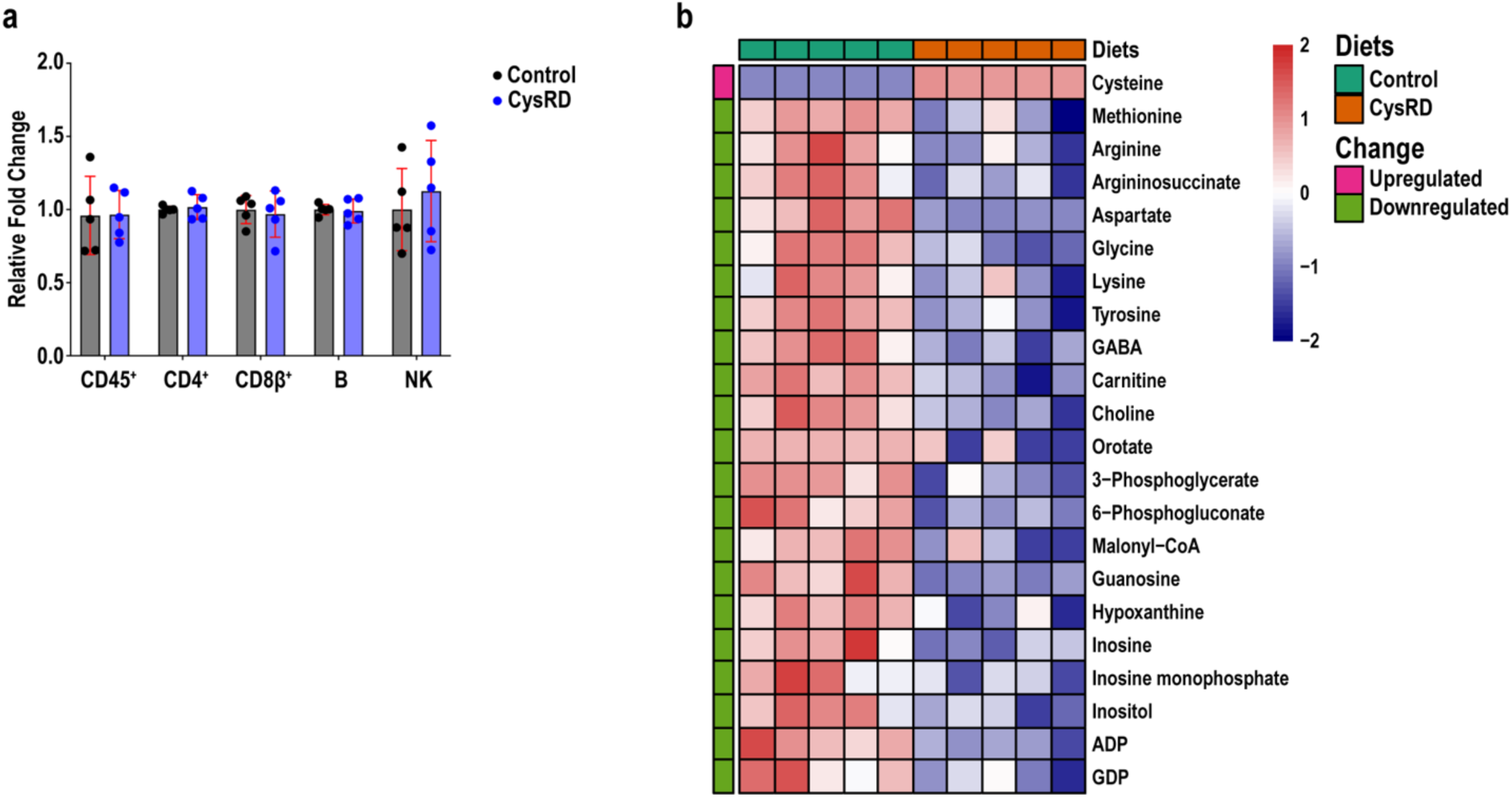
Cysteine-rich diet in the colon does not affect crypt-associated CD8αβ^+^ T cells or CoA generation. **a**, Relative fold changes of colon intraepithelial lymphocytes from control- and CysRD-fed mice. **b**, Heatmap with most significant metabolites affected by CysRD in the colon. n = 5 biological replicates. Unpaired two-tailed Student’s t-tests (**a**,**b**). Data are mean ± s.d.

## Notes

### Competing Interest Statement

The authors have declared no competing interest.

